# Applying Self-Supervised Tissue Patch Encoders to Enable Cellular Graph Encoding

**DOI:** 10.1101/2025.06.10.658651

**Authors:** Zhiyuan Ding, Alex Baras

**Affiliations:** Department of Health Sciences Informatics, Johns Hopkins University; Department of Pathology, Johns Hopkins University

## Abstract

Recent advances in computation pathology have seen the development of various forms of foundational models that have enabled high-quality, general-purpose feature extraction from tissue patches. However, most of these models are somewhat limited in their ability to capture cell-to-cell spatial relationships essential for understanding tissue microenvironments. To address this issue, self-supervised learning (SSL) applied to cellular graph models is a promising avenue of investigation as this would afford one the ability to capture cell-level interpretability of the tissue microenvironment and learn relationships from the cellular topology. Compared to SSL on tissue patches, developing cellular graph models requires considerable computation and algorithmic effort, starting with accurate cell detection/segmentation from tissue and then subsequently requiring adequate cell-level feature extraction. In this study, we demonstrate that **a pretrained tissue patch encoder, originally trained in the context of the token used in conventional vision transformer frameworks can also serve effectively as a cellular graph encoder for cell-based patches**. We report three key findings: 1) the pretrained tissue patch encoder acts as a powerful graph-level feature extractor for cellular graphs in tissue patches; 2) it also can function as a celllevel feature extractor, allowing each cell to be contextualized within its local tissue microenvironment environment; 3) the cell patch size and positional embeddings are critical factors for high-quality cellular graph feature extraction. To validate and support these insights, we evaluate our approach on various tissue region classification tasks from three different types of cancers. Experimental results demonstrate that our method achieves performance comparable to the current state-of-the-art foundation models in computational pathology. These results highlight the potential of pretrained tissue patch encoders in cellular graph modeling, which can enable insights into how the cellular features and topology are related to various biological processes and phenomena.

## 1 Introduction

Self-supervised learning (SSL) has enabled the development of a series of high-quality feature extractors specifically tailored for tissue patches in computational pathology (CPath). These extractors have achieved promising results across tasks on different levels and have shown robust capabilities in various contexts [11, 30, 10]. Their structures rely exclusively on vision transformers (ViT) [16] that operate at the image patch level or above, segmenting images into patches based on regular grids, encoding localized texture features, and aggregating these into image-level representations through self-attention mechanisms. Despite the strong performance on patch and slide-level tasks, their ability to capture finegrained cell-to-cell relationships remains constrained. In [11], while the fine-tuned model demonstrates superior performance compared to other competitive approaches, its effectiveness in cell-level segmentation falls short of the high accuracy achieved in patch or slide level downstream tasks. This limitation can be attributed to the strategies of current SSL frameworks, such as MAE [22], MoCo v2 [12], and DINO v2 [41], which focus primarily on predicting masked patches or enforcing feature invariance across augmented views, thereby emphasizing localized or holistic features rather than inter-cellular interactions, a feature rather specific to CPath.

As an alternative modeling strategy, instead of using images of tissue regions, cellular graphs are constructed where vertices represent individual detected cells or nuclei. This approach enables direct representation of cell-to-cell interactions through vertex connections in the graph structure. Therefore, developing SSL frameworks for cellular graphs is appealing, as they offer an effective means of extracting tissue features while capturing cell-level interpretability and learning directly from the cellular topology [7, 43, 38, 32, 25, 46, 33]. However, several challenges currently limit the implementation of cellular graph SSL. First, constructing cellular graphs requires reliable cell detection and segmentation, which can be resource-intensive. Even when accurate segmentation is achieved, segmenting all cells within whole slide images for large-scale datasets is costly. For example, slides in the Non-Small Cell Lung Cancer (NSCLC) dataset [13, 24], which will be used in this study, typically contain around 400,000 detected nuclei. Second, cellular graph construction requires robust cell-level feature extraction, but no standardized pipeline exists for defining node features across datasets. Consequently, SSL frameworks trained on specific cell-level features may not generalize well to different cellular feature extraction pipelines. Additionally, cellular graphs exhibit considerable variation in cell counts across different tissue regions. For example, in the Prostate cANcer graDe Assessment (PANDA) [8] dataset used in our experiments, each graph typically contains fewer than 70 detected nuclei on average, but over 10% graphs contain more than 300 nuclei. This broad range of cell counts presents challenges: graphs with fewer cells are susceptible to over-smoothing issues [9] with increased hidden layers, while those with many cells risk over-squashing issues [50], where critical details may be compressed or lost. These factors make training an SSL model based on cellular graphs a relatively inefficient task compared to conventional tissue image-based methods.

In this study, we pose the following question: **is it necessary to train an SSL model from scratch to obtain a cellular graph encoder?** We find that the answer is **no**. A pretrained tissue patch encoder, originally trained with token-based structures in conventional ViT frameworks, can effectively function as a cellular graph encoder for cell-based patches. Our approach redefines the concept of tokens by using cell patches as tokens rather than traditional (nonoverlapping) image patches, allowing us to encode tissue structure through cellular graphs based on these cell tokens instead of relying on whole-image representations. We systematically examine factors influencing the efficacy of this cellular graph encoder strategy and observe that, with an optimized cell patch size and well-calibrated positional embeddings, the adapted cellular graph encoder achieves performance comparable to or even outperforms conventional image-based encoders. With this cell-centric encoding strategy, distinct and informative cell-level features can also be extracted. These enriched celllevel features can be seamlessly integrated into other multi-instance learning (MIL) models and graph neural networks (GNNs). Overall, the key contributions are as follows:

- A novel cellular graph encoder is introduced, leveraging a pretrained tissue patch image encoder to extract high-quality features at both the graph and cell levels, providing a versatile foundation for various tissue analysis tasks.
- Systematic studies conducted on this cellular graph encoder reveal that the size of cell patches and positional embeddings are two critical factors that significantly impact the quality of feature extraction.
- Extensive experiments are conducted on three tissue-level classification datasets (prostate, breast, and lung), with results that strongly demonstrate the effectiveness, interpretability, and generalization ability of the proposed method across different tissue contexts.

## 2 Related Work

### SSL models in Cpath

Recently, with the availability of large-scale CPath datasets, SSL has been increasingly used to extract task-agnostic feature representations. Pretrained SSL models generally fall into three categories: (1) vision-only foundation models [10, 11, 55, 53], which derive visual features using contrastive, reconstruction, or hybrid approaches; (2) vision-language models [35, 3], where images are aligned with text data to enhance the contextualization of visual features; and (3) other multi-modal models [1, 29, 28], which extract modality-agnostic features by aligning images with other modalities or materials. While some recent studies, such as [46], have explored SSL on cellular graphs, as we discuss in Section 1, this approach proves less effective than conventional methods and requires additional information during training, such as assuming cellular graphs from one slide are similar to each other. In our work, we do not conduct any SSL training directly. Instead, we treat cellular graphs as a structural variant of standard tissue images, with a cellular-centric organization, and investigate how to utilize SSL models trained on conventional tissue images to encode cellular graphs.

### Cellular graphs

Research has demonstrated a connection between cell topology and various cancer types [20, 34, 56]. Cellular graphs, constructed on individual cells, directly model inter-cellular interactions along edges through message-passing neural networks [19], allowing predictions to be explicitly based on cell interactions. Prior work on cellular graphs can generally be grouped by task type: (1) cell-level classification tasks [32, 33], which identify cell subtypes based on each cell’s intrinsic properties and its surrounding microenvironment, and (2) tissue or slide-level tasks [60, 43, 38], which leverage single-graph or multiple-graph features for region-of-interest (ROI) or slide-level prediction. Our study centers on the tissue-level task, where image patch or cell-level information is aggregated without employing hierarchical structures. This allows us to directly compare the efficacy of conventional image encoding strategy with our cellular graph-based encoding approach. From the model perspective, we also observe that several cellular graph-based models utilize transformer or graph transformer architectures for cellular graph encoding [38, 43, 32, 59]. Typically, these models rely on pre-extracted cell-level features, which are obtained either through separate feature extraction networks or by using handcrafted feature sets. In contrast, our approach directly employs SSL ViTs pretrained on pathological images to perform both cell-level and graph-level feature extraction, thereby bypassing the need for separated cell feature extraction and aggregation steps.

### Utilizing spatial information for tissue-level CPath analysis

In CPath, spatial information is typically incorporated at various levels due to the large number of instances involved. A prevalent paradigm for instance-level feature aggregation is MIL [14, 27, 36, 45, 48, 57], in which instances are aggregated by applying weighted averaging based on instance-level attention scores. This method minimally uses spatial information, as each instance is only influenced by others in the aggregation phase, without explicitly encoding spatial relationships between instances. GNNs [32, 33, 60,43, 38, 31, 49, 42], on the other hand, utilize spatial information to different extents depending on the structure of their adjacency matrices and edge attributes. When using only a discrete adjacency matrix, where vertices are either connected or not, spatial encoding is restricted to the fundamental topology of vertex connections. Some GNN models extend spatial encoding by incorporating detailed edge attributes, such as precise vertex distances and inter-cellular texture features, achieving a more granular representation of spatial relationships. Transformers[38, 43, 32, 59, 54] leverage spatial information similar to GNNs with enhanced spatial encoding. Positional embeddings are used to capture the relative positions between patches, enabling a detailed spatial encoding. In this study, we adopt the transformer framework for cellular graph encoding and assess its performance against models that encode spatial information at varying levels of detail / spatial resolution.

## 3 Method

We introduce our framework for encoding cellular graphs using a pretrained ViT to achieve high-quality graph and cell-level feature extraction (see Fig. 1). The motivations and preliminary are discussed in Section 3.1, while details of the proposed cellular graph encoder are provided in Section 3.2. An analysis of influential factors is presented in Section 3.3.

**Figure 1:**
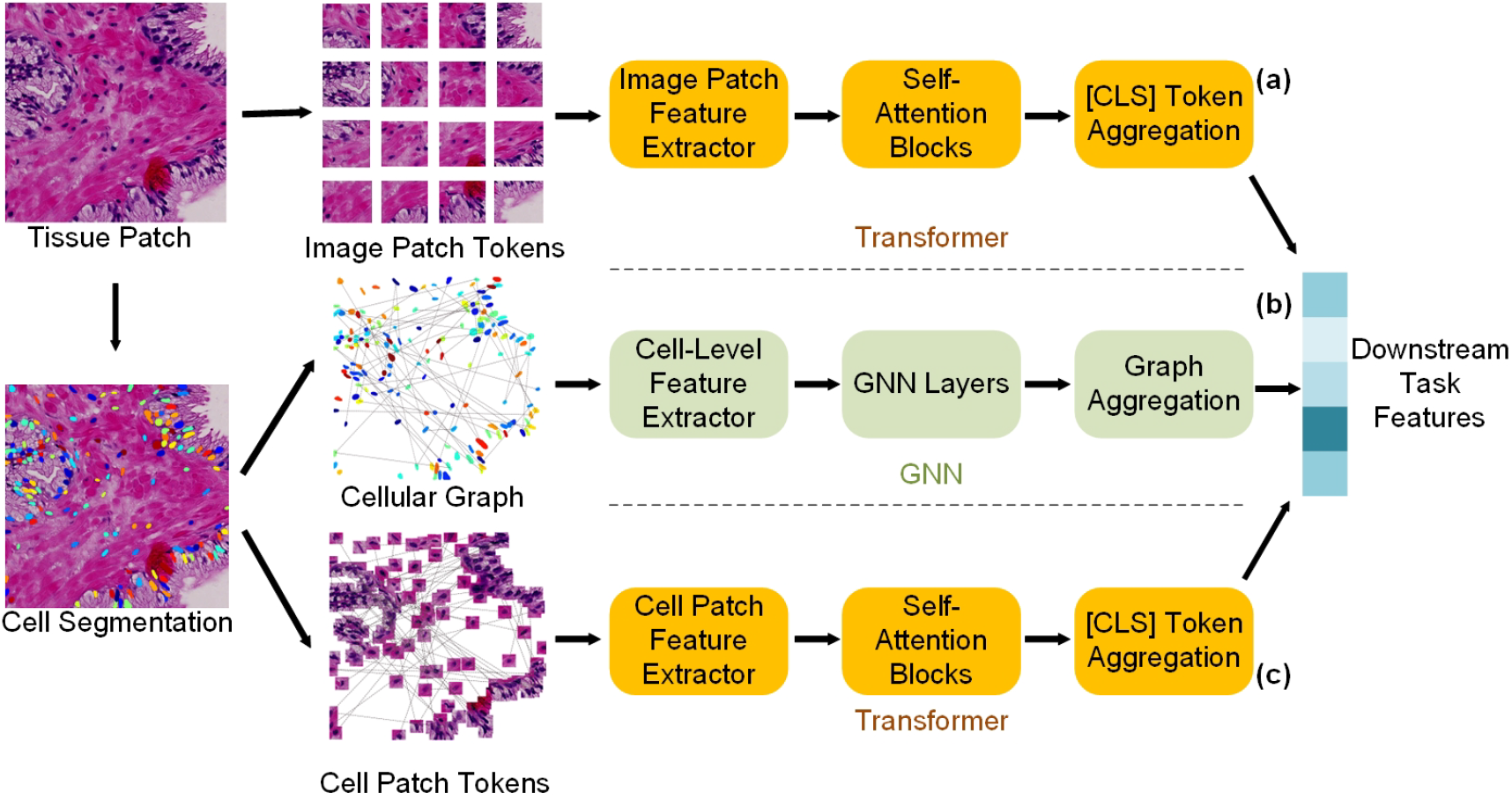
Feature extraction frameworks: (a) SSL pretrained ViT for tissue images; (b) GNN-based cellular graphs; (c) proposed SSL-pretrained ViT adapted for cellular graphs. The same color represents the same model structure and weights.

### 3.1 Motivations and Preliminary

#### Motivations

In CPath, tissue patches can be encoded using two distinct strategies: (1) treating images as conventional computer vision inputs, where local patches are defined on regular grids, and interactions between biological entities are implicitly captured by expanding receptive fields in deeper network layers; (2) using a cellular-centric approach, which focuses only on regions containing detected cells or nuclei for further analysis. With the availability of large-scale histopathology datasets, the feature extraction quality of approach (1) has achieved promising results across various CPath tasks. Conversely, GNNs for cellular graphs are not yet widely adapted with SSL strategies to generate robust, task-agnostic features. Given that the cellular graph is constructed from regions containing detected cells, as in highthroughput microscopy [5], if assuming each local image patch contains exactly one cell, a ViT can directly serve as the encoder, functioning identically to a cellular graph encoder without further processing. We further extend this idea to handle irregular cell localizations by introducing cell patch tokens for cellular graph encoding.

#### Feature extraction paradigms for tissue patches and cellular graphs

For a conventional ViT, given a tissue patch image 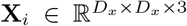, **X**_*i*_ is divided into local image patches 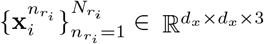 for feature extraction, with a total number of local patches 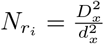. The ViT *f*_*V*_ then encodes the local patches 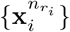 into a feature representation 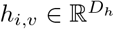. In cellular graphs, however, cells or nuclei are first identified by cell segmentation models, after which regions containing the detected cells 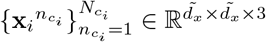 are used for single-cell level feature extraction. In this study, we use StarDist [47] for nuclei segmentation. Previous cellular graph analysis models generally follow a two-step paradigm. First, single-cell level feature extractors *f*_*SC*_, such as pretrained ResNet [23] / transformer on ImageNet, and handcrafted feature extractor[37], extract singlecell feature 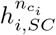 for each cell patch 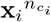. Then, GNN layers *f*_*MP*_, such as GraphSAGE [21], Graph Attention Network (GAT) [52], and Graph Isomorphic Network (GIN) [58], propagate features across vertices and their neighbors. Finally, a graph-level aggregation layer *f*_*Aggr*_ combines node-level features into a graph representation.

### 3.2 Cellular Graph Encoder

#### Cellular graph GNNs are functionally identical to ViT

As depicted in Fig. 1, cellular graph feature extraction and ViT-based tissue image feature extraction can both be understood as three-step processes. In the first step, features are extracted at the single-cell or image patch level: cellular graphs utilize cell-level feature extractors *f*_*SC*_, while ViTs use patch embedding layers (or also with several layers in self-attention blocks) to transform image patches into tokens. In the second step, features are contextualized based on their relationships with neighboring elements: this is achieved through GNN layers *f*_*MP*_ in cellular graphs, and self-attention mechanisms in ViTs. Finally, an aggregation operation gathers the contextualized features into a global representation—graph-level aggregation layers, such as mean or sum pooling, are used in cellular graphs, while the [CLS] token in ViTs aggregates information across all tokens. These parallel processes illustrate that GNN encoders in cellular graphs function analogously to ViT encoders, both transforming localized features into representations that capture various forms of spatial information.

#### Encoding cellular graph through pretrained ViT

Based on the observation above, cellular graphs, represented by cellular patches 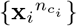 and corresponding spatial information, can be directly encoded using a pretrained ViT to obtain high-quality features at both the node and graph levels (see Fig. 1 (c)). We use weights from [11] in this research. In particular, cell patch 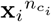 is resized to 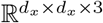 for processing with the ViT, and convolutional layers with the following self-attention blocks work as *f*_*SC*_ to extract features at the single-cell level. To enable self-attention-based encoding between cells, proper positional embeddings are required for the cells. In a regular grid-based ViT, the positional embedding for a patch located at (*x, y*) is determined by projecting (*x, y*) into 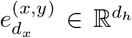 by the embedding layer and adding it to the patch’s token as input for the self-attention blocks. As illustrated in Fig. 2 (a), to compute the positional embeddings for each cell, we first interpolate the positional embeddings 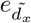 for the regular grid patches of size 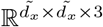, as proposed in [16], to avoid issues with cells detected outside the bounds of the current coordinate system. As in the figure, the blue star represents a detected nucleus, which cannot be encoded using the original coordinate system (orange dashed line), but can be encoded using the interpolated coordinate system (blue dashed line). Subsequently, the cell-level positional embedding 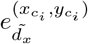 is computed using bilinear interpolation, considering the four nearest grid centroids, as indicated by the red star in the figure. This interpolated cell-level positional embedding is then added to the features extracted from the cell patches, and the remaining process follows the standard inference procedure for regular grid image patches (use selfattention blocks as *f*_*MP*_ in GNNs). At the graph level, features are directly obtained from the [CLS] token. For cell-level features, we collect the output after the convolutional layer as well as the output tokens from the self-attention blocks.

**Figure 2:**
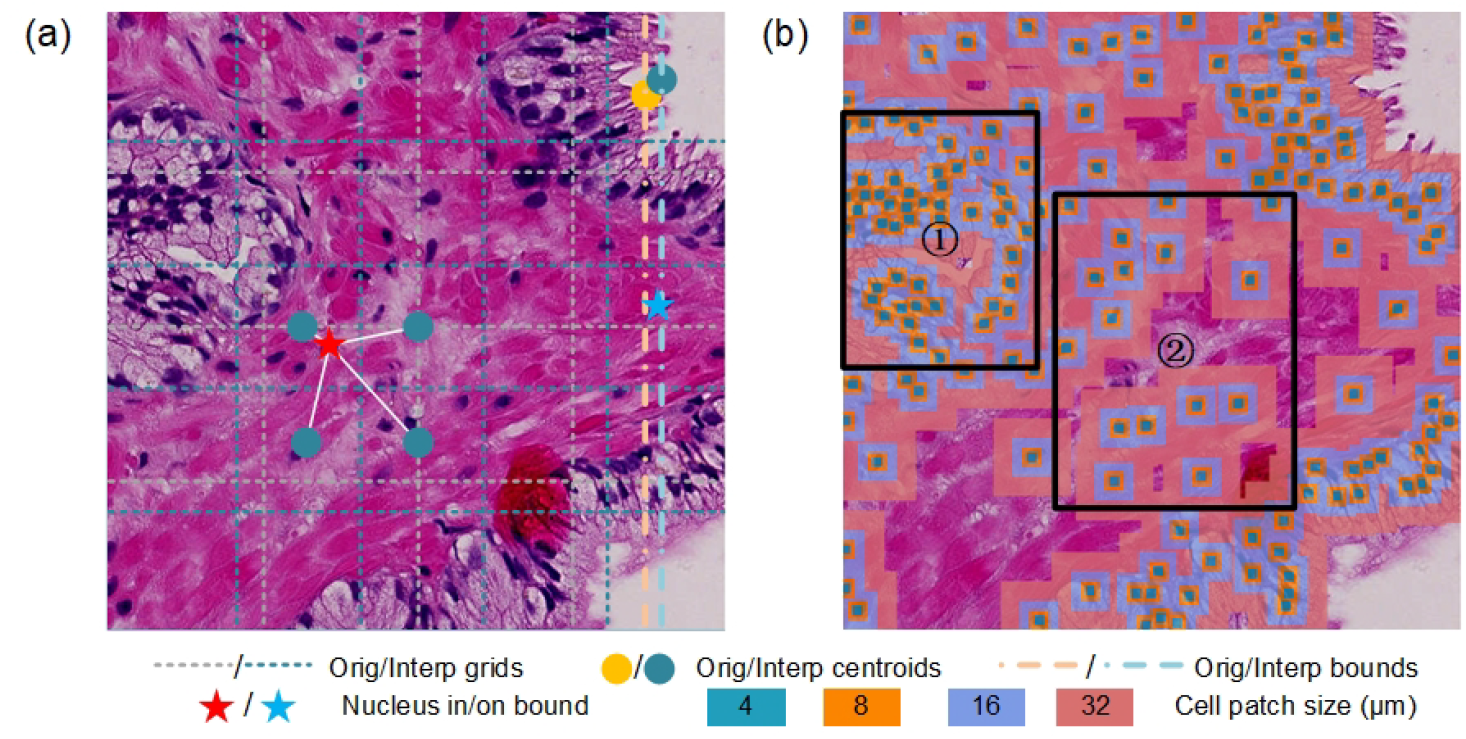
(a) Example of cell positional embedding interpolation process. Original and interpreted results are presented. Grids, centroids, and bounds are image patch boundaries, centroids, and maximal positional encoding ranges, respectively. Stars are examples of positions to encode. (b) **Covering ranges of cellular graphs with varied cell patch sizes**. Different colors represent different cell patch sizes in *µ*m. ① is an example of epithelial gland region and ② is stroma.

### 3.3 Key Factors That Influence Performance

We examine the influence of various factors that could potentially affect the prediction performance of the features, and identify two factors that have significant impacts on the results.

#### Spatial information of cells

Introducing spatial information directly encodes the cell topology into the extracted features. In contrast, without spatial information, node features are contextualized in the weakest possible way, similar to MIL, where each instance only implicitly understands the existence of other instances through self-attention. If the cell topology contributes to the prediction, then incorporating spatial information will enhance performance. **Covering ranges of cell patches**. Enlarging the cell-level patch gradually increases the microenvironment information of the cells, thereby improving its prediction ability. To illustrate this concept, we visualize the covering range of cells with varying sizes, as shown in Fig. 2 (b) for the prostate tissue patch in (a). When the cell size is 4 *µ*m, only parts of the nuclei are contained in the cellular patch, and the resulting cellular graph primarily reflects the nuclei morphology. As the cell size increases to 8 *µ*m, epithelial gland structure (cell clusters, an example is the left bounding box) can be captured, so the obtained cellular graph starts to encode the dense cell cluster information. Moreover, when the cell patch size grows to 16 and 32 *µ*m, the patches cover increasingly larger regions of the tissue, include more details of the epithelial glands, and start encoding the microenvironment of the stroma (spread of cells, an example is the right bounding box). However, we also observe that the performance does not continue to increase indefinitely with respect to the cell patch sizes. This upper bound is related to feature redundancy and is determined by the specific tissue types.

### 3.4 Downstream Task Evaluation

Both graph and node level features are extracted for cellular graphs using a pretrained ViT [11], which was trained on over 100 million in-house histopathology H&E images. At the graph level, features with and without positional embeddings are extracted and evaluated by training multilayer perceptrons (MLPs) with varied cell patch sizes. At the node level, features from convolutional layers and self-attention blocks, both with and without positional embeddings, are extracted and further encoded through attention-based MIL (ABMIL) [27], GraphSAGE [21], GAT [52], and GIN [58] to assess the effectiveness of feature extraction quality. We also evaluate the extracted feature in an unsupervised manner through optimal transport. More details can be found in **Supplemental**.

## 4 Experiment and Results

### 4.1 Dataset

To verify the effectiveness of our proposed cellular graph encoding strategy, we conducted experiments directly on tissue-level tasks. This strategy allows for a direct performance comparison between tissue images and their cellular graph representations while minimizing potential confounding factors, such as hierarchical structures required for feature aggregation in slide-level tasks and the influence of nuclei segmentation quality on cell-level classification tasks. In this study, we developed and evaluated three specific tissue-level tasks to rigorously assess the proposed encoding strategy’s performance in pathology analysis. For further details on preprocessing steps and dataset description, please refer to the **Supplemental**.

#### Prostate cancer tissue level grading

The PANDA challenge dataset [8] provides a large-scale collection of 10,616 digitized prostate biopsies, labeled according to the International Society of Urological Pathology (ISUP) or Gleason grading (GG) system for sample-level prediction. In this study, we focus on a subset of samples from Radboud University, specifically selected for their pixel-level annotations. To standardize the dataset for our purposes, we transform these annotations into tissue patch-level labels consistent with the slide-level grading groups defined in [51, 18, 44]. Unlike the original slide-level grouping, we analyze fixed-size tissue patches sampled from the biopsies. This approach yields 493,928 tissue images of 6 classes from 5,049 biopsy slides. In 4.2 and 4.3, we further split the task into cancer detection (which in theory should be relatively independent of local cell topology) and cancer grading (predicting from GG1 to GG5, which should depend more in cell topology in this medical context based on our current understand) to observe the effect of incorporating spatial information.

#### Breast Invasive Carcinoma (BRCA) subtyping

The Cancer Genome Atlas (TCGA) BRCA dataset [24, 2] includes 954 histopathology slides, comprising 787 from patients with Infiltrating Ductal Carcinoma (IDC) and 167 from patients with Infiltrating Lobular Carcinoma (ILC). Directly cropping these whole-slide images into tissue patches would result in an overwhelmingly large number of patches, many of which would not contain relevant cancerous tissue. To address this issue, we applied a tumor region detection network [15], trained on the CAMELYON16 dataset [4], to filter out patches with a higher likelihood of containing tumor regions. By this approach, we effectively reduced the number of patches to those with substantial tumor presence. For this study, we selected a subset of 200 slides (100 per class), ultimately obtaining 65,907 tissue patches. Since these filtered patches represent tumor regions, we can directly use the slide-level labels for tissue-level subtyping.

#### Non-Small Cell Lung Cancer Subtyping

The TCGA NSCLC dataset [24, 39] comprises 1,120 histopathology slides, including samples from 530 Lung Adenocarcinoma (LUAD) and 490 Lung Squamous Cell Carcinoma (LUSC) patients. Following a similar approach to the BRCA dataset, we applied the tumor region detection network [15] to identify tumor regions in a subset of 200 slides. This preprocessing step yielded a total of 70,551 tissue patches, with slide-level labels directly assigned to each patch. **General experiment standards**. Due to the large number of patches in the datasets, we do not employ a cross-validation strategy. Instead, we split each dataset once into training, validation, and test sets at the slide level to ensure consistent data partitioning across all experiments. The ratio of train/validation/test samples at the slide level is 3:1:1. For performance evaluation, we report the area under the curve (AUC), accuracy, and F1 score, reporting test performance for the model with the lowest validation loss (as is the convention). For further details on experimental settings, please refer to the **Supplemental**.

### 4.2 Graph-level Performance

We first evaluate the proposed graph-level cellular graph features directly encoded by the pretrained ViT. Specifically, we assess the impact of positional embedding and cell patch sizes by training MLPs with features generated under each condition. The performance across these four tasks is summarized in Fig. 3. For the detailed metrics of the performance, please refer to the **Supplemental**.

**Figure 3:**
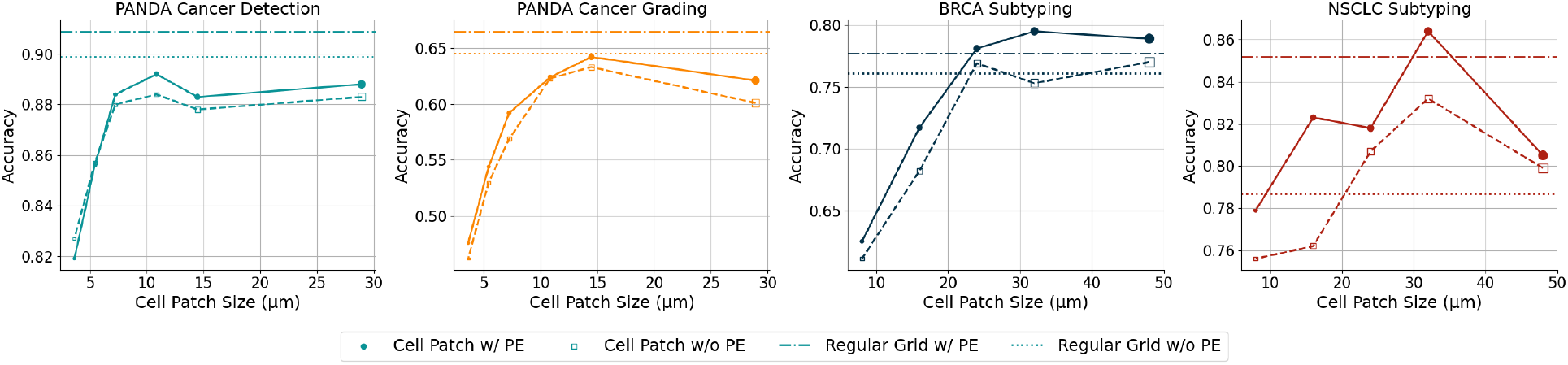
Comparison of cellular graph level feature performances across different cell patch sizes, with and without positional embeddings, on multiple tasks. ViT features extracted using tissue images serve as the reference with or without positional embedding.

Cellular graph features achieve comparable performance to conventional tissue image-based features across all tasks. Specifically, while cellular graphs perform slightly lower in the two PANDA dataset tasks, they show improved results on the BRCA and NSCLC subtyping tasks. With appropriately chosen cell sizes, cellular graphs outperform direct tissue image encoding, achieving a +1.8% increase in accuracy for BRCA subtyping and a +1.2% increase for NSCLC subtyping.

Integrating positional encoding enhances the quality of extracted features for both cellular graph and tissue image representations in nearly all cases. For cellular graph-level features, the addition of positional embedding yields an average accuracy improvement of 1.4% increase for the PANDA grading task, 2.8% for BRCA subtyping, and 3.1% for NSCLC subtyping.

Increasing the cell patch size generally enhances prediction performance until reaching a threshold where performance gains become saturated. Specifically, for the PANDA cancer detection task, accuracy improves by more than 6.5% as the cell patch size increases from 4 to 12 *µ*m. Similarly, for the PANDA cancer grading task, accuracy improves by 14%, and for BRCA subtyping, by 17% as the cell patch size increases from 8 to 32 *µ*m, while for NSCLC subtyping, accuracy improves by 8.5%. Saturation is observed across all tasks except for PANDA cancer detection. This phenomenon is likely due to the homogeneity of features among cell patches, where features from different positions become increasingly similar at larger patch sizes. Additionally, prostate cancer detection, unlike grading, can often be achieved by observing nuclear morphology alone [17, 26]. This is also supported by the relatively stable performance (82% accuracy) observed even at the smallest cell patch size of 4 *µ*m, with which only the nuclei morphology information can be obtained for each cell. Thus, choosing a task and data specific cell patch size can meaningfully enhance feature extraction and predictive performance.

### 4.3 Node-level Performance

To verify that pretrained SSL models can also extract informative node-level features, we follow classic CPath frameworks, specifically ABMIL and GNN, to assess node-level feature quality. For the GNN model, we employ GAT v2 [6] with edge attributes, as it performs slightly better than alternative models, though results are generally stable across different GNN models and settings. We use the output tokens of self-attention blocks without positional embedding as the node feature to ensure the least spatial information is used. In constructing the cellular graph, nodes are connected using a k-nearest neighbors (kNN) approach with *k* = 5, excluding connections between nodes separated by more than 30 *µ*m, a commonly used configuration for cellular graph construction.

As illustrated in Fig. 4, using cell patches with an appropriate cell patch size yields improved performance compared to ABMIL applied to conventional tissue patch features. The relationship between cell patch size and performance mirrors the pattern observed at the graph level: increasing cell patch size initially boosts prediction performance until it reaches saturation. For GNN models, increasing the cell patch size improves accuracy by 7% for PANDA cancer detection, 8% for PANDA cancer grading, 14.6% for BRCA subtyping, and 8.5% for NSCLC subtyping. When comparing ABMIL and GNN performance, we observe that incorporating spatial information enhances predictive capability, aligning with our findings for graph-level features. Specifically, comparing peak performance for GNN and ABMIL, spatial information contributes a 0.5% improvement in PANDA cancer identification, 0.9% in PANDA cancer grading, 1.8% in BRCA subtyping, and 1.3% in NSCLC subtyping.

**Figure 4:**
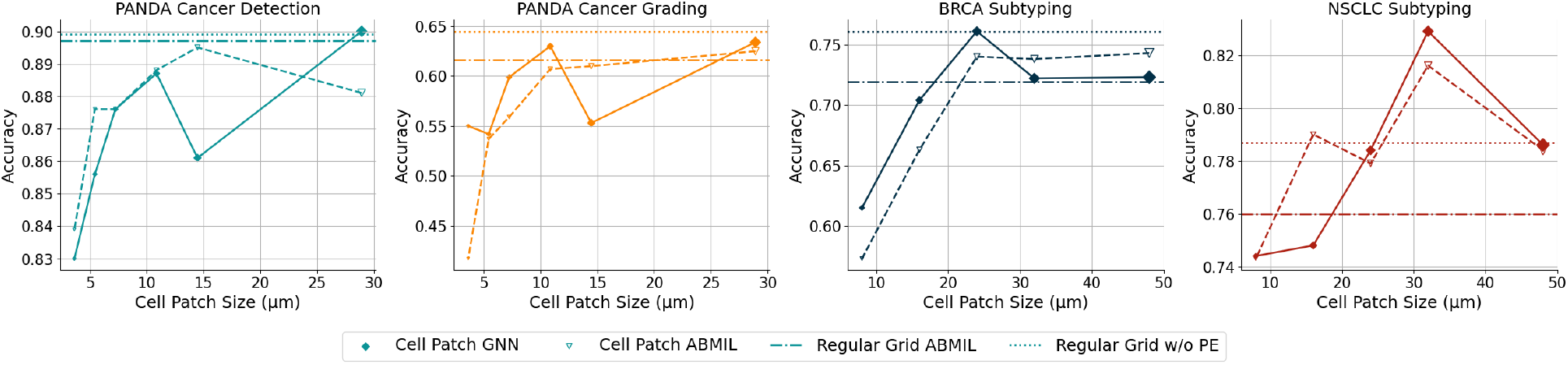
Comparison of cellular node-level features across different cell patch sizes, with and without positional embeddings, on multiple tasks. ViT features without positional embeddings and ABMIL with tissue images patch tokens (regular grid) serves as the reference.

### 4.4 Other Settings and Interpretation

#### Impact of GNN structure

To assess the generalization capability of our proposed cellular graph encoding method and demonstrate that the specific GNN structure is not the primary factor, we evaluate various GNN architectures, with results presented in Table 1. Although the ViT-based encoding strategy achieves the highest performance, differences between encoding models are relatively minor. Specifically, when compared to the best GNN model, altering the model structure results in accuracy decrease most of 3.3% for the PANDA subtyping task, 4.5% for the BRCA subtyping task, and 2.1% for the NSCLC subtyping task. We also investigated additional GNN structures that may impact predictive performance and details are provided in the **Supplemental**.

**Table 1:**
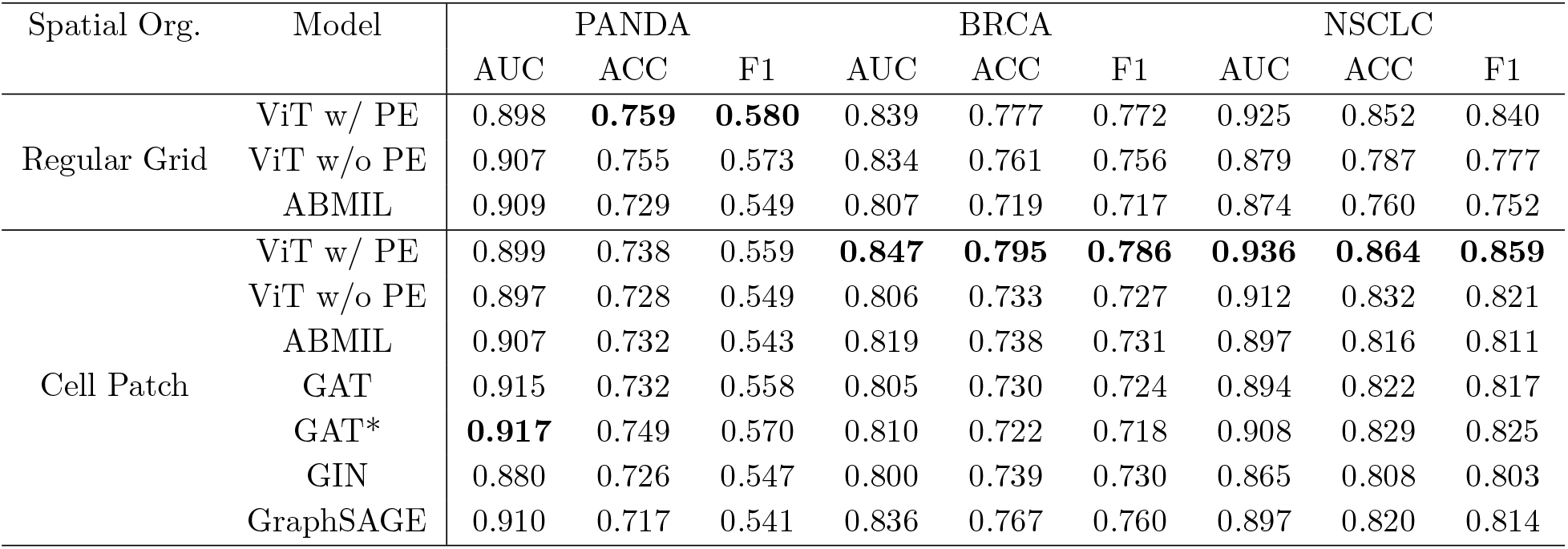
Comparison of graph-level and node-level features across various encoders. ‘Spatial Org.’ refers to different spatial organization strategies applied to tissue images (tissue image) and cellular graphs (cell patch). GAT* denotes a GAT with edge attributes specified by distance values. The optimal cell patch size is selected for each dataset.

#### Saliency map based interpretation

The saliency maps (Fig. 5) show that in general higher values (red color) are seen in the malignant epithelial cells as compared to background stroma and normal epithelial cells. When comparing the the saliency maps of the regular grid approach (as in used with ViT) to the cell patch approach, one can begin to associated certain topologies of well formed circular glands in GG1 to the the poorly formed glands (if at all) in higher grades such as GG4 and GG5. We also conduct a quantitative analysis of the saliency map in the **Supplemental**.

**Figure 5:**
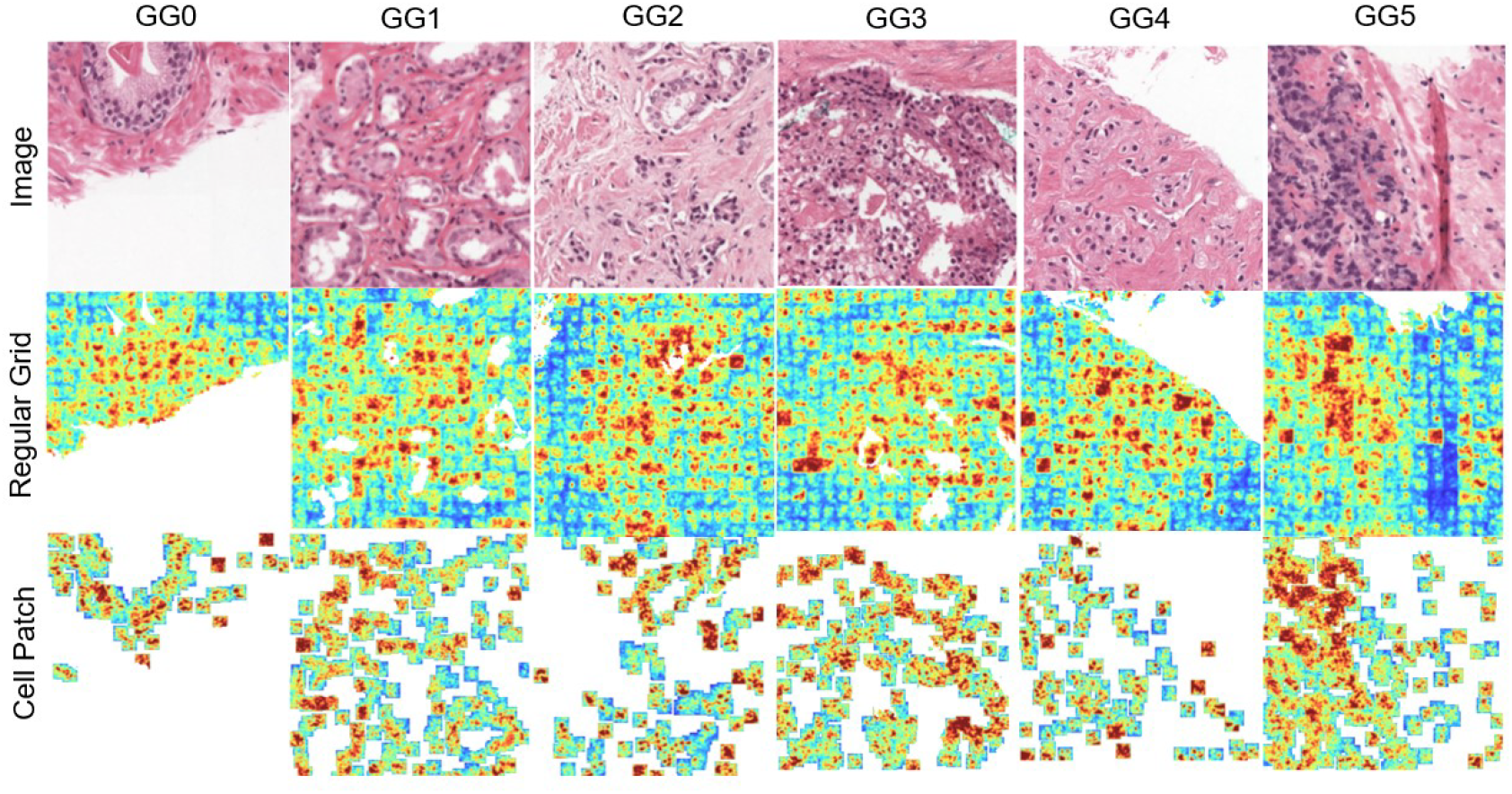
Saliency map comparison. Results are shown for the PANDA dataset. Each column represents a grade, with the first-row tissue images, the second-row saliency map of regular grid encoding, and the third-row representing cell patch encoding.

## 5 Conclusion

In this paper, we introduce a method to encode cellular graphs using pretrained SSL models, specifically trained on tissue images, for cell-centric histopathology analysis. We evaluate our framework across three distinct tissue-level classification tasks, covering varied tissue types (prostate, breast, and lung). The proposed cellular encoding strategy achieves comparable performance to whole-tissue image encoding and surpasses it when optimal cell patch size and spatial information are incorporated at both the graph and node levels. This work contributes to the field of cellular graph analysis by enabling high-quality feature extraction and analysis through pretrained models on tissue images. Future work will explore encoding cellular graphs for multiplex images, which incorporate multiple biomarkers within a single section [40]. Additionally, applying the proposed framework to other task types, such as cell classification and slide classification, may yield promising and interpretable results for histological analysis.

