## Supplementary materials for "Applying Self-Supervised Tissue Patch Encoders to Enable Cellular Graph Encoding"

### Supplementary Material for: Applying Self-Supervised Tissue Patch Encoders to Enable Cellular Graph Encoding

#### Appendix

In this appendix, we provide additional details and further analyses not covered in the manuscript. Section A describes the preprocessing steps, including tissue patch collection, cell segmentation, feature extraction, and cell graph construction. Section B details the three datasets used in our experiments. Section C outlines the network structures, training configurations, and visualization settings. Section D expands on the experimental results, including graph-level classification, node-level classification performance, the layers used for node feature extraction, and evaluations of other cellular graph encoding strategies. In Section E, we systematically analyze the feature distributions in an unsupervised manner using two distributional distance metrics, demonstrating performance trends consistent with predictive ability. Finally, Section F presents a quantitative evaluation of the saliency maps.

#### A. Preprocessing

Our preprocessing pipeline consists of four main steps to derive both node-level and graph-level features: tissue patch filtering, cell segmentation, feature extraction, and cellular graph construction.

**Tissue patch filtering.** In this study, we use a patch size of 512 with a step size of 256. Following this, the tissue tile filtering process takes all tiles from this sliding window approach as input to yield a filtered output. For the PANDA dataset, we use a subset of biopsies from Radboud University, which includes pixel-level annotations. Given that these annotations are relatively noisy, we first generate a scaled mask at a lower resolution, which serves as the tile-level ground truth. Using this mask, we categorize tissue components into background, benign tissues (Subtypes 1 & 2), and cancerous tissues (Subtypes 3, 4, and 5). Tiles containing more than 50% background are excluded from further analysis. For the BRCA and NSCLC datasets, where pixel-level annotations are unavailable, we employ a tumor region detection network [5]. This network, a U-Net model trained to predict pixel-level probabilities of tumor regions, is used to filter out tumor-related regions for the subtyping task. Specifically, we utilize pretrained weights from the CAMELYON16 challenge [3], a breast cancer H&E slide classification task with pixel-level annotations, to predict tumor regions. Tissue patches with an average tumor probability exceeding 0.1 are selected for subsequent subtyping analysis.

**Cell segmentation.** We use StarDist [16] for nuclei detection, leveraging a U-Net model trained to predict both probability maps and geometric polygons for each nucleus. To enable efficient cell segmentation on tissue images, we reimplement the StarDist cell segmentation pipeline to process batches of tissue images, resulting in a time savings of approximately 60% compared to the default StarDist API. However, a bottleneck remains in the non-maximum suppression (NMS) step, which we intend to address in future work. For cell segmentation, we utilize pretrained network weights from the MoNuSeg challenge [9] and the TNBC dataset [13]. We set a detection threshold of 0.3 and an overlap ratio of 0.3 for NMS across all three tasks. A pathologist reviews the segmentation results to ensure accuracy, enabling reliable performance in downstream analysis.

**Feature extraction.** In this study, we use the ViT model with pretrained weights from [4] as a feature extractor. For tissue images, we resize the input from 512 to 256 pixels for feature extraction, resulting in $16 \times 16$ local image patches. This resizing is close to the SSL training input size of 224 and facilitates comparison with the default cell patch size of 32 used in this study. For the cellular graph, we first select nuclei within the tissue image and use corresponding cell patches with relative spatial locations as input for the ViT. Each cell patch is resized to $16 \times 16$, matching the default local patch size used in [4]. Positional embeddings for each cell are then computed as described in the Method section. We collect representations with and without positional embeddings for analysis of image or graph-level features. We extract features from both a convolutional layer and the final self-attention block's output tokens for node-level analysis, omitting positional embeddings.

**Cellular graph construction.** Using the extracted features, cellular graphs can be represented in two ways: (1) as a 1024-dimensional graph-level feature derived from the

[CLS] token of the ViT, or (2) as a cellular graph where each node has a 1024-dimensional attribute. For the cellular graphs, we define edges between nodes based on the k-nearest neighbor strategy, a widely used approach in cellular graph construction. Specifically, we set $k = 5$, and cellular distances exceeding 60 pixels (about 30 $\mu$m) are excluded from the analysis. Cellular graphs with fewer than six cells are also removed from further analysis.

#### B. Details for Datasets

**PANDA dataset.** Following the ISUP grading system, we define tissue patch and cellular graph-level labels as described in prior studies [6, 15, 17]. Specifically, for each sample, we consider the proportions of different tissue types within a patch, denoted as: $s_1$ for background tissue, $s_2$ for normal tissue, $s_3$ for cancerous subtype 3, $s_4$ for cancerous subtype 4, and $s_5$ for cancerous subtype 5.

From these proportions, we derive several key ratios to characterize the tissue composition. The foreground ratio, defined as

$$r_1 = \frac{s_3 + s_4 + s_5}{s_2 + s_3 + s_4 + s_5},$$

represents the proportion of cancerous tissue relative to the total non-background area. Additionally, we calculate the relative contributions of each cancerous subtype with:

$$r_2 = \frac{s_3}{s_3 + s_4 + s_5}, \quad r_3 = \frac{s_4}{s_3 + s_4 + s_5}, \quad r_4 = \frac{s_5}{s_3 + s_4 + s_5}.$$

The graph-level labels are then assigned based on these calculated ratios, as Table 1 summarizes. These labels facilitate consistent characterization of tissue composition across samples. A summary of sample statistics for datasets processed using these criteria is provided in Table 2.

**BRCA dataset.** After the preprocessing steps, a total of 179 samples remain following tumor detection. The slide-level label is then assigned as the tissue image label for subtyping. Statistics for the BRCA dataset are presented in Table 3.

**NSCLC dataset.** After the preprocessing steps, a total of 182 samples remain following tumor detection. The slide-level label is then assigned as the tissue image label for subtyping. Statistics for the NSCLC dataset are presented in Table 4.

| Class | Definition |
|---|---|
| GG0 | $r_1 = 0$ |
| GG1 | $r_1 > 0.05$ & $r_2 = 1$ |
| GG2 | $r_1 > 0.05$ & $r_4 < 0.05$ & $r_2 > r_3$ |
| GG3 | $r_1 > 0.05$ & $r_4 < 0.05$ & $r_3 > r_2$ |
| GG4 | $r_1 > 0.05$ & $r_3 = 1$<br>$r_1 > 0.05$ & $r_4 > 0.05$ & $r_2 > 0.05$ & $r_3 < r_2 < r_4$<br>$r_1 > 0.05$ & $r_4 > 0.05$ & $r_2 > 0.05$ & $r_3 < r_4 < r_2$<br>$r_1 > 0.05$ & $r_4 > 0.05$ & $r_2 > 0.05$ & $r_4 < r_3 < r_2$ |
| GG5 | $r_1 > 0.05$ & $r_2 < 0.05$ & $r_4 < 0.05$<br>$r_1 > 0.05$ & $r_4 > 0.05$ & $r_2 > 0.05$ & $r_4 < r_2 < r_3$<br>$r_1 > 0.05$ & $r_4 > 0.05$ & $r_2 > 0.05$ & $r_2 < r_3 < r_4$<br>$r_1 > 0.05$ & $r_4 > 0.05$ & $r_2 > 0.05$ & $r_2 < r_4 < r_3$ |

Table 1. **Tissue image and graph level label definition for PANDA dataset.** GG0 is normal tissue, and GG1 to GG5 follows the definition for ISUP score.

| Class/Split | Train | Validation | Test |
|---|---|---|---|
| GG0 | 164293 | 53041 | 55440 |
| GG1 | 41004 | 13985 | 13969 |
| GG2 | 55725 | 18681 | 18407 |
| GG3 | 19659 | 7228 | 6068 |
| GG4 | 9917 | 2992 | 3560 |
| GG5 | 6006 | 1902 | 2121 |

Table 2. **Statistics of PANDA dataset.**

| Set/Class | ILC | IDC |
|---|---|---|
| Slide | 82 | 97 |
| Train | 16774 | 21479 |
| Validation | 6555 | 6575 |
| Test | 5522 | 9002 |

Table 3. **Statistics of BRCA dataset.**

| Set/Class | LUAD | LUSC |
|---|---|---|
| Slide | 88 | 94 |
| Train | 18338 | 25779 |
| Validation | 8047 | 3929 |
| Test | 8191 | 6267 |

Table 4. **Statistics of NSCLC dataset.**

#### C. Network and General Settings

**Graph-level feature downstream task settings.** For graph-level tasks, MLPs are employed for downstream analysis. Specifically, a 3-layer MLP with a hidden dimension of 128 is used across all graph-level tasks, incorporating dropout with a rate of 0.4 and ReLU activations between layers. A final softmax layer is applied for output classification.

**Node-level feature downstream task settings.** For AB-MIL, the instance-level feature extractor is a 3-layer MLP with dropout set to 0.4 and ReLU activation. The attention mechanism is then learned from the extracted features through a 2-layer MLP. Following a weighted average operation, a linear layer with softmax activation is used to generate the final prediction. For GNNs, we utilize different message-passing layers based on the chosen GNN types, with the number of message-passing layers set to 3. ReLU activation and dropout at a rate of 0.4 are applied between each GNN layer. We evaluate various types of node and graph aggregation methods; please refer to Table D for details. After graph-level aggregation, a linear layer is used to produce the final prediction.

**General Training Settings** The network is trained with an initial learning rate of $5\times10^{-4}$ using the Adam optimizer. A batch size of 256 is used for all experiments. Cross-entropy loss is employed, with class weights applied to address the class imbalance. Specifically, these weights are determined by the inverse ratio of sample counts in the training set. Based on validation loss tolerance, early stopping is implemented with patience of 10 epochs for node-level tasks and 20 epochs for graph-level tasks.

**Saliency Map** The saliency map is generated by computing the gradient of the image with respect to the ground truth class. For visualization, we use the absolute values of the gradient, applying min-max normalization with clipping at a tolerance range from 0.025 (and 1-0.025) to 0.05 (and 1-0.05) quantiles. Additionally, background regions and holes are removed in the visualization to prevent unwanted artifacts in the analysis.

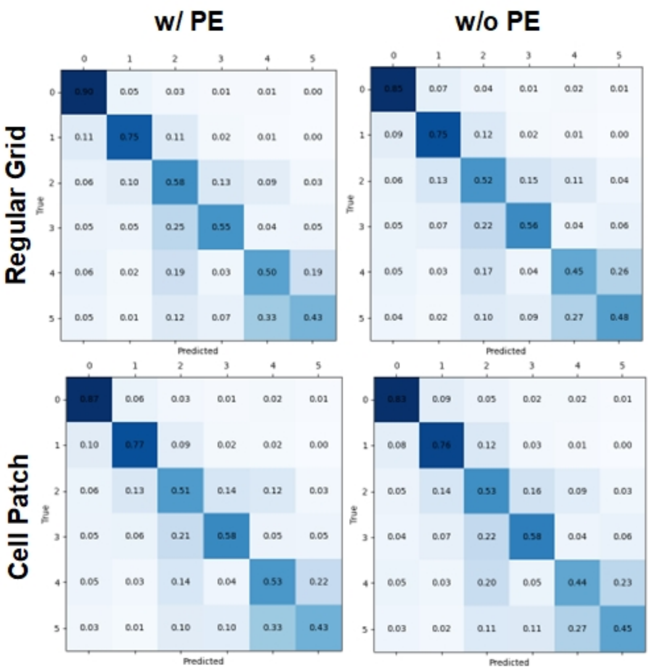


Figure 1. **Confusion matrix of PANDA cancer prediction task.** Results from regular grid and cell patch based ViTs with or without positional encoding.

#### D. Details of results.

**Graph level Results.** Results for the graph-level experiments are presented in Table 5. Specifically, we observe that the proposed ViT-based cellular graph encoding approach achieves performance comparable to that of the conventional tissue image-based ViT. Regarding cell patch size, performance improves as the cell patch size increases, stabilizing once the patch size reaches a sufficient threshold. Analyzing the effects of positional embedding, we find that incorporating spatial information generally improves results, except for AUC in the PANDA dataset. This observation aligns with the findings discussed in the manuscript. Interestingly, the best AUC for the PANDA dataset is achieved using a regular grid without positional embedding, suggesting that spatial information may be less critical than initially anticipated. Furthermore, a detailed comparison of confusion matrices for regular grid-based and cell patch-based predictions (16 $\mu$m resolution) is presented in Fig. 1. The results reveal that cell patch-based encoding demonstrates superior performance in grading cases with GG 3, 4, and 5, while regular grid-based encoding outperforms in identifying cases with GG 0, 1, and 2.

**Node level results.** Results of node-level features are presented in Table 6. We observe that the incorporation of spatial information enhances predictive performance, with the GNN achieving the highest performance across all three tasks. Additionally, increasing the cell patch size progressively boosts performance, though the optimal patch size varies across datasets, with each dataset exhibiting a different saturation point.

**Comparison of different aggregation strategies.** In this section, we outline the network structures examined in this study. In addition to the results presented in the manuscript, we also evaluated several commonly used GNN structures within cell-level and patch-level CPath GNN frameworks. Specifically, we investigated the effects of various node aggregation strategies, graph aggregation strategies, the incorporation of residual connections between GNN layers [10], and the use of jumping knowledge [18]. For these experiments, the optimal cell patch sizes were selected: 16 $\mu$m for the PANDA dataset and 32 $\mu$m for both the BRCA and NSCLC datasets. Results are provided in Table 7. We observe that none of these strategies significantly impact the final performance.

**Comparison of node feature extraction layer.** We also evaluate the performance of various feature extraction approaches, with the results summarized in Table 8. The findings indicate that node-level features derived from self-attention blocks significantly outperform those extracted solely after the convolutional block, suggesting that self-attention blocks function not only as feature extractors but also as aggregators of neighborhood information.

| PE options + Spatial Org. | cell patch size ($\mu$m) | PANDA | | | BRCA | | | NSCLC | | |
|---|---|---|---|---|---|---|---|---|---|---|
| | | AUC | ACC | F1 | AUC | ACC | F1 | AUC | ACC | F1 |
| w/ PE + Regular Grid | - | 0.898 | **0.759** | **0.580** | 0.839 | 0.777 | 0.772 | 0.925 | 0.852 | 0.840 |
| w/o PE + Regular Grid | - | **0.907** | 0.755 | 0.573 | 0.834 | 0.761 | 0.756 | 0.879 | 0.787 | 0.777 |
| w/ PE + Cell Patch | 4 | 0.849 | 0.648 | 0.438 | - | - | - | - | - | - |
| | 6 | 0.878 | 0.682 | 0.486 | - | - | - | - | - | - |
| | 8 | 0.889 | 0.713 | 0.526 | 0.667 | 0.625 | 0.618 | 0.859 | 0.779 | 0.764 |
| | 12 | 0.898 | 0.744 | 0.561 | - | - | - | - | - | - |
| | 16 | 0.899 | 0.738 | 0.559 | 0.792 | 0.717 | 0.712 | 0.890 | 0.823 | 0.816 |
| | 24 | - | - | - | 0.839 | 0.781 | 0.771 | 0.913 | 0.818 | 0.810 |
| | 32 | 0.898 | 0.725 | 0.546 | **0.847** | **0.795** | **0.786** | **0.936** | **0.864** | **0.859** |
| | 48 | - | - | - | 0.845 | 0.789 | 0.781 | 0.881 | 0.805 | 0.792 |
| w/o PE + Cell Patch | 4 | 0.849 | 0.648 | 0.438 | - | - | - | - | - | - |
| | 6 | 0.866 | 0.681 | 0.481 | - | - | - | - | - | - |
| | 8 | 0.889 | 0.702 | 0.516 | 0.636 | 0.611 | 0.595 | 0.842 | 0.756 | 0.755 |
| | 12 | 0.904 | 0.726 | 0.549 | - | - | - | - | - | - |
| | 16 | 0.897 | 0.728 | 0.549 | 0.737 | 0.682 | 0.673 | 0.880 | 0.762 | 0.758 |
| | 24 | - | - | - | 0.821 | 0.769 | 0.757 | 0.905 | 0.807 | 0.801 |
| | 32 | 0.896 | 0.712 | 0.535 | 0.806 | 0.733 | 0.727 | 0.912 | 0.832 | 0.821 |
| | 48 | - | - | - | 0.817 | 0.770 | 0.759 | 0.883 | 0.799 | 0.784 |

Table 5. **Performance comparison of cell patch size for graph level features.** PE represents positional embeddings. Spatial Org. is spatial organization, which includes regular grid (conventional ViT) and Cell Patch based ViT.

| Spatial Org. + Model | cell patch size ($\mu$m) | PANDA | | | BRCA | | | NSCLC | | |
|---|---|---|---|---|---|---|---|---|---|---|
| | | AUC | ACC | F1 | AUC | ACC | F1 | AUC | ACC | F1 |
| Regular Grid ABMIL | - | 0.909 | 0.729 | 0.549 | 0.807 | 0.719 | 0.717 | 0.874 | 0.760 | 0.752 |
| Cell Patch + ABMIL | 4 | 0.852 | 0.640 | 0.435 | - | - | - | - | - | - |
| | 6 | 0.875 | 0.682 | 0.485 | - | - | - | - | - | - |
| | 8 | 0.893 | 0.696 | 0.506 | 0.620 | 0.573 | 0.570 | 0.838 | 0.743 | 0.741 |
| | 12 | 0.903 | 0.728 | 0.542 | - | - | - | - | - | - |
| | 16 | 0.899 | 0.907 | 0.532 | 0.741 | 0.663 | 0.659 | 0.857 | 0.790 | 0.772 |
| | 24 | - | - | - | 0.797 | 0.740 | 0.729 | 0.876 | 0.779 | 0.773 |
| | 32 | 0.898 | 0.728 | 0.541 | 0.819 | 0.738 | 0.731 | 0.897 | 0.816 | 0.811 |
| | 48 | - | - | - | 0.818 | 0.743 | 0.734 | 0.878 | 0.784 | 0.776 |
| Cell Patch + GNN | 4 | 0.864 | 0.663 | 0.465 | - | - | - | - | - | - |
| | 6 | 0.888 | 0.696 | 0.499 | - | - | - | - | - | - |
| | 8 | 0.904 | 0.709 | 0.521 | 0.614 | 0.615 | 0.592 | 0.840 | 0.744 | 0.743 |
| | 12 | 0.916 | 0.745 | 0.559 | - | - | - | - | - | - |
| | 16 | **0.917** | **0.749** | **0.570** | 0.764 | 0.704 | 0.695 | 0.876 | 0.748 | 0.744 |
| | 24 | - | - | - | 0.828 | **0.761** | **0.751** | 0.886 | 0.784 | 0.777 |
| | 32 | 0.908 | 0.724 | 0.550 | 0.810 | 0.722 | 0.718 | **0.908** | **0.829** | **0.825** |
| | 48 | - | - | - | **0.833** | 0.723 | 0.722 | 0.872 | 0.786 | 0.778 |

Table 6. **Performance comparison of cell patch size for node level features.** Spatial Org. is the spatial organization, which includes regular grid (conventional ViT) and Cell Patch based ViT. Here the utilized GNN model is GAT v2 with edge attribute defined by distances between nodes.

| Spatial Org. | Model | PANDA | | | BRCA | | | NSCLC | | |
|---|---|---|---|---|---|---|---|---|---|---|
| | | AUC | ACC | F1 | AUC | ACC | F1 | AUC | ACC | F1 |
| Regular Grid | ViT w/ PE | 0.898 | **0.759** | **0.580** | 0.839 | 0.777 | 0.772 | 0.925 | 0.852 | 0.840 |
| | ViT w/o PE | 0.907 | 0.755 | 0.573 | 0.834 | 0.761 | 0.756 | 0.879 | 0.787 | 0.777 |
| | ABMIL | 0.909 | 0.729 | 0.549 | 0.807 | 0.719 | 0.717 | 0.874 | 0.760 | 0.752 |
| Cell Patch | ViT w/ PE | 0.899 | 0.738 | 0.559 | **0.847** | **0.795** | **0.786** | **0.936** | **0.864** | **0.859** |
| | ViT w/o PE | 0.897 | 0.728 | 0.549 | 0.806 | 0.733 | 0.727 | 0.912 | 0.832 | 0.821 |
| | ABMIL | 0.907 | 0.732 | 0.543 | 0.819 | 0.738 | 0.731 | 0.897 | 0.816 | 0.811 |
| | GAT | 0.915 | 0.732 | 0.558 | 0.805 | 0.730 | 0.724 | 0.894 | 0.822 | 0.817 |
| | GAT* | 0.917 | 0.749 | 0.570 | 0.810 | 0.722 | 0.718 | 0.908 | 0.829 | 0.825 |
| | GIN | 0.880 | 0.726 | 0.547 | 0.800 | 0.739 | 0.730 | 0.865 | 0.808 | 0.803 |
| | GraphSAGE | 0.910 | 0.717 | 0.541 | 0.836 | 0.767 | 0.760 | 0.897 | 0.820 | 0.814 |
| | GNN + max node aggr. | 0.913 | 0.723 | 0.542 | 0.804 | 0.723 | 0.717 | 0.910 | 0.829 | 0.820 |
| | GNN + max graph aggr. | 0.901 | 0.732 | 0.532 | 0.808 | 0.732 | 0.727 | 0.895 | 0.819 | 0.814 |
| | GNN + sum graph aggr. | 0.744 | 0.693 | 0.407 | 0.791 | 0.761 | 0.756 | 0.813 | 0.793 | 0.789 |
| | GNN + residual | **0.920** | 0.740 | 0.557 | 0.811 | 0.745 | 0.735 | 0.909 | 0.828 | 0.823 |
| | GNN + jumping knowledge | 0.912 | 0.732 | 0.553 | 0.827 | 0.756 | 0.751 | 0.892 | 0.820 | 0.813 |

Table 7. **Comparison of graph-level and node-level features across various encoders.** 'Spatial Org.' refers to different spatial organization strategies applied to tissue images (tissue image) and cellular graphs (cell patch). GAT* denotes a GAT with edge attributes specified by distance values. The optimal cell patch size is selected for each dataset. To verify the impact of node aggregation and graph aggregation strategies, we use GraSAGE as the basic network structure. For residual and jumping knowledge, GAT* is used as basic network structure.

| Feature | PANDA | | | BRCA | | | NSCLC | | |
|---|---|---|---|---|---|---|---|---|---|
| | AUC | ACC | F1 | AUC | ACC | F1 | AUC | ACC | F1 |
| SA | **0.917** | **0.749** | **0.570** | **0.810** | **0.772** | **0.778** | **0.908** | **0.829** | **0.825** |
| Conv | 0.867 | 0.673 | 0.476 | 0.612 | 0.657 | 0.625 | 0.608 | 0.638 | 0.587 |

Table 8. **Comparison of feature extraction position.** GATv2 with edge attribute is used as the encoder. SA represents the feature extracted after self-attention blocks and Conv represents the feature extracted only after the convolutional block. No positional embedding is used for node level feature extraction process.

#### E. Feature Distribution Analysis

In addition to evaluating prediction performance, we examine feature distributions under varying conditions to assess the impact of cell patch sizes and the incorporation of positional embeddings. Qualitatively, feature distributions are visualized using UMAP [11], while quantitatively, distributional distances are measured between the gold standard (features extracted using a regular grid-based ViT) and those obtained with our proposed cell patch-based ViT approach.

**Distribution Descriptors.** In this paper, we introduce a methodology employing two descriptors to quantify distributional gaps between feature spaces, aiming to elucidate unsupervised similarities and differences. Specifically, we leverage the Gromov-Wasserstein Distances (GWD) [12, 14] and Wasserstein Distances (WD) as metrics, computed in both bottom-up and top-down approaches, to measure the distributional divergence between two domains. Formally, as in the Method section, let the extracted features from the source and target domains be denoted as $H_v^s$ and $H_v^t$, respectively, where $H_v^j = \{h_{i,v}^j | i = 1, 2, \dots, N_j\}$ represents the feature set belonging to domain $j$, with $j \in \{s, t\}$. The GWD is designed to compute the minimal distance $D_{GW}$ between the two domains, formulated as:

$$\begin{aligned} D_{GW} = \min_{\mathbf{T}} & \sum_{k,l,p,q} L(\mathbf{C}_{s,k,l}, \mathbf{C}_{t,p,q}) \mathbf{T}_{k,l} \mathbf{T}_{p,q}, \\ \text{s.t. } & \mathbf{T1} = \mathbf{p}_s, \\ & \mathbf{1T} = \mathbf{p}_t, \\ & \mathbf{T} \geq 0. \end{aligned} \quad (1)$$

Here, $\mathbf{C}j \in \mathbb{R}^{Nj \times N_j}$ represents the pairwise cost matrix derived from $H_v^j$, for which we use the Euclidean distance in this study. The optimization target, $\mathbf{T} \in \mathbb{R}^{N_j \times N_j}$, is a transport matrix that aligns the distributions of the source and target domains. This transport matrix satisfies constraints ensuring its row and column sums correspond to the respective probability distributions of the source and target domains while maintaining non-negativity. Direct computation of GWD involves calculating the cost matrix and optimizing the transport matrix of size $N_j \times N_j$, which is computationally infeasible in our case. To address this limitation, we propose two alternative approaches. The first approach adopts a bottom-up strategy, operating on batches of samples to approximate the computation. The second approach models the distributions of each domain, fitting parametric representations to these distributions, and subsequently using the learned models as descriptors for calculating distributional distances.

**Bottom-up distributional distance.** We compute the GWD for batches of samples and average across batches as the domain-level distributional distance. Formally, with a given batch size $B$, the bottom-up GWD is formulated as:

$$D_{bottom-up} = \frac{1}{N_j} \sum_{n_b=1}^{N_j/B} D_{GW}(H^s_{n_b,v}, H^t_{n_b,v}), \quad (2)$$

where $H^j_{n_b,v} = \{h^j_{i,v} | i = \frac{N}{B} \times (n_b - 1) + 1, \frac{N}{B} \times (n_b - 1) + 2, \ldots, \frac{N}{B} \times n_b\}$, is the $n_b$-th batch of samples. In this computation process, we assume a uniform distribution of samples in both the source and target domains, as the sampling is performed uniformly.

**Up-bottom distributional distance.** In this up-bottom approach, we first employ a parametric model to estimate the feature distributions of the domains. The distributional distances are then calculated between these parametric models to approximate the domain-level distributional differences. Formally, we fit Gaussian Mixture Models (GMM) to represent the domain distributions:

$$\mathbf{p}_j = \sum_{c_j} \pi_{c_j} \mathcal{N}(\mu_{c_j}, \Sigma_{c_j}), \quad (3)$$

where the nonnegative mixture coefficient $\sum_{c_j} \pi_{c_j} = 1$, $\mathcal{N}$ is Gaussian distribution with mean $\mu_{c_j} \in \mathbb{R}^{N_j}$ and covariance $\Sigma_{c_j} \in \mathbb{R}^{N_j \times N_j}$. The WD between two Gaussians is formulated in previous researches [7, 8]:

$$D_{WG}(G_1, G_2) = ||\mu_1 - \mu_2||^2 + tr(\Sigma_1 + \Sigma_2 - 2\Sigma_2^{\frac{1}{2}} \Sigma_1 \Sigma_2^{\frac{1}{2}})^{\frac{1}{2}}. \quad (4)$$

We follow the idea from [2], and approximate the distributional distances as a barycenter optimal transport problem:

$$D_{up-bottom} = \sum_{c_s, c_t} \pi_{c_s} \pi_{c_t} D_{WG}(G_{c_s}, G_{c_t}). \quad (5)$$

**Results.** For both visualization and distributional distance analyses, we employ the Euclidean distance metric to quantify feature space differences. Experiments are conducted on the PANDA dataset, leveraging its extensive sample size, with 20% of the dataset used for the subsequent analysis. For $D_{bottom-up}$, batch size 256 is used. For $D_{up-bottom}$, GMM is assumed to have 10 kernels.

The visualization results are presented in Fig. 2 and Fig. 3. These results demonstrate that as the cell patch size increases, the feature distribution progressively aligns more closely with the feature distribution of a regular grid. When the cell patch size is smaller than 12 $\mu$m, the feature space 'collapses', resulting in significantly lower predictive performance compared to features with larger cell patch sizes, which exhibit feature spaces more similar to a regular grid. Interestingly, we also observe a 'collapse' in the feature space for cell patch sizes of 32 $\mu$m. This behavior is likely attributable to the increased homogeneity of cell patches as the patch size continues to grow. Since direct observation of UMAP-based feature distributions poses challenges for systematic and objective comparison, we compute distributional distances as evaluation metrics. The quantitative result is shown in Fig. 4. Specifically, we use features extracted from a regular grid ViT with positional embeddings as the reference standard and compute the distributional distances of other features relative to it. A larger distributional distance indicates greater dissimilarity between the features of the two domains. The results reveal an anticorrelated relationship between distributional distance and predictive performance as the cell patch size changes. When positional embeddings are included, the learned features exhibit greater similarity to the reference standard. This observation aligns with the findings presented in the manuscript, emphasizing that both cell patch size and positional embeddings are critical factors influencing the quality of the encoded features.

#### F. Saliency Map Analysis

To quantitatively assess the differences between saliency maps learned by different models, we follow the methodology outlined in [1] and compute the Structural Similarity Index Measure (SSIM) and Spearman Rank Correlation (SRC) as indicators of saliency map distances. To eliminate the influence of uncovered regions in the cell patch encoding method, these metrics are calculated only within the cell patch regions, rather than across the entire image. The results, presented in Fig. 5, show that the saliency map distances do not align with the criteria observed for predictive performance or feature distributional distances. This discrepancy suggests that models trained with cell patches and regular grids may approach the problem using fundamentally different patterns of learned representation and demonstrates the necessity for developing cellular graph based analytical tools for interpretable analytical model.

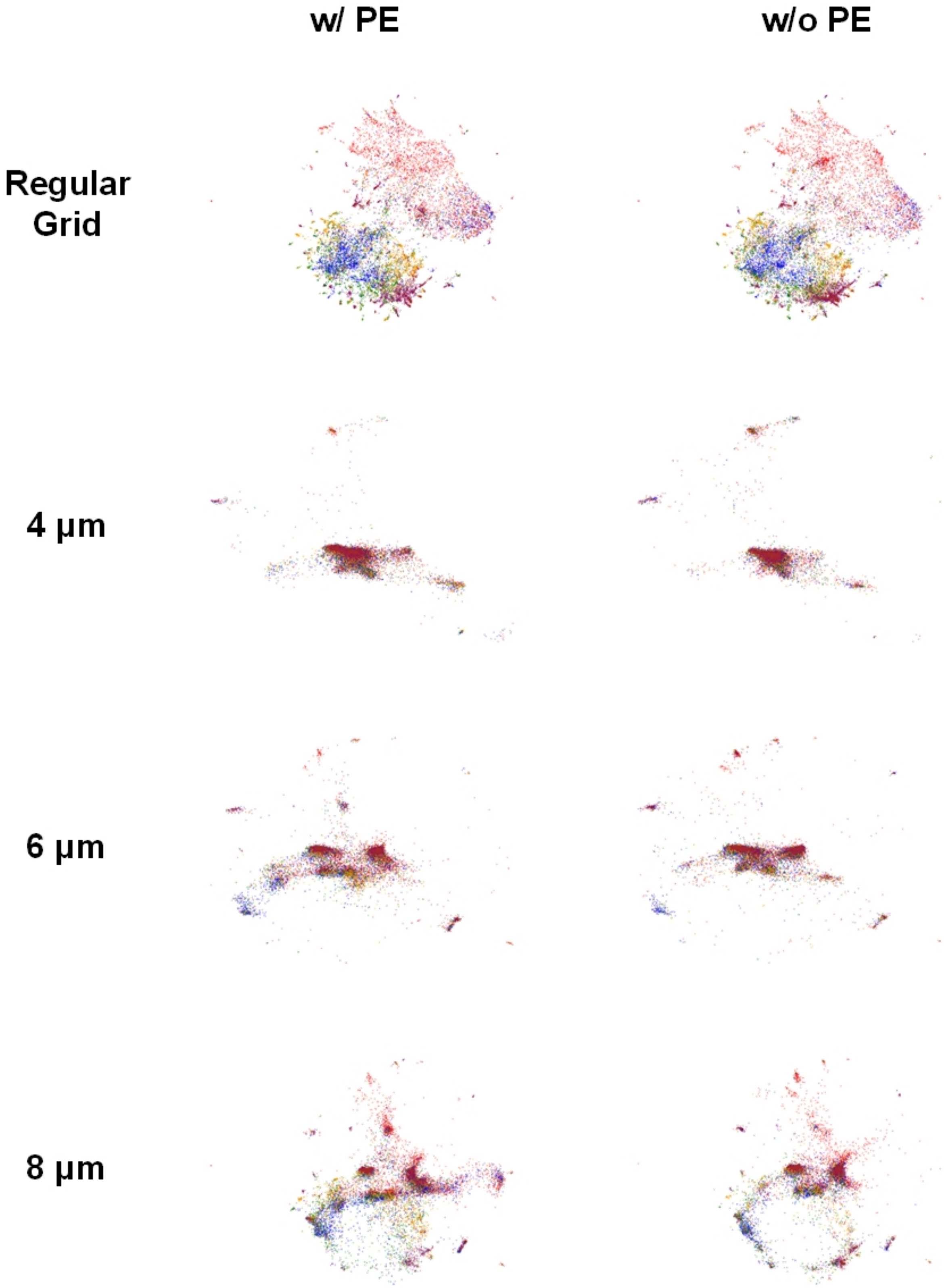


Figure 2. **Feature distributions.** Regular grid represents features extracted from conventional ViT. Both with and without positional embeddings are checked. Different colors represent different grading. Cell patch sizes 4,6,8 $\mu$m are included in this figure.

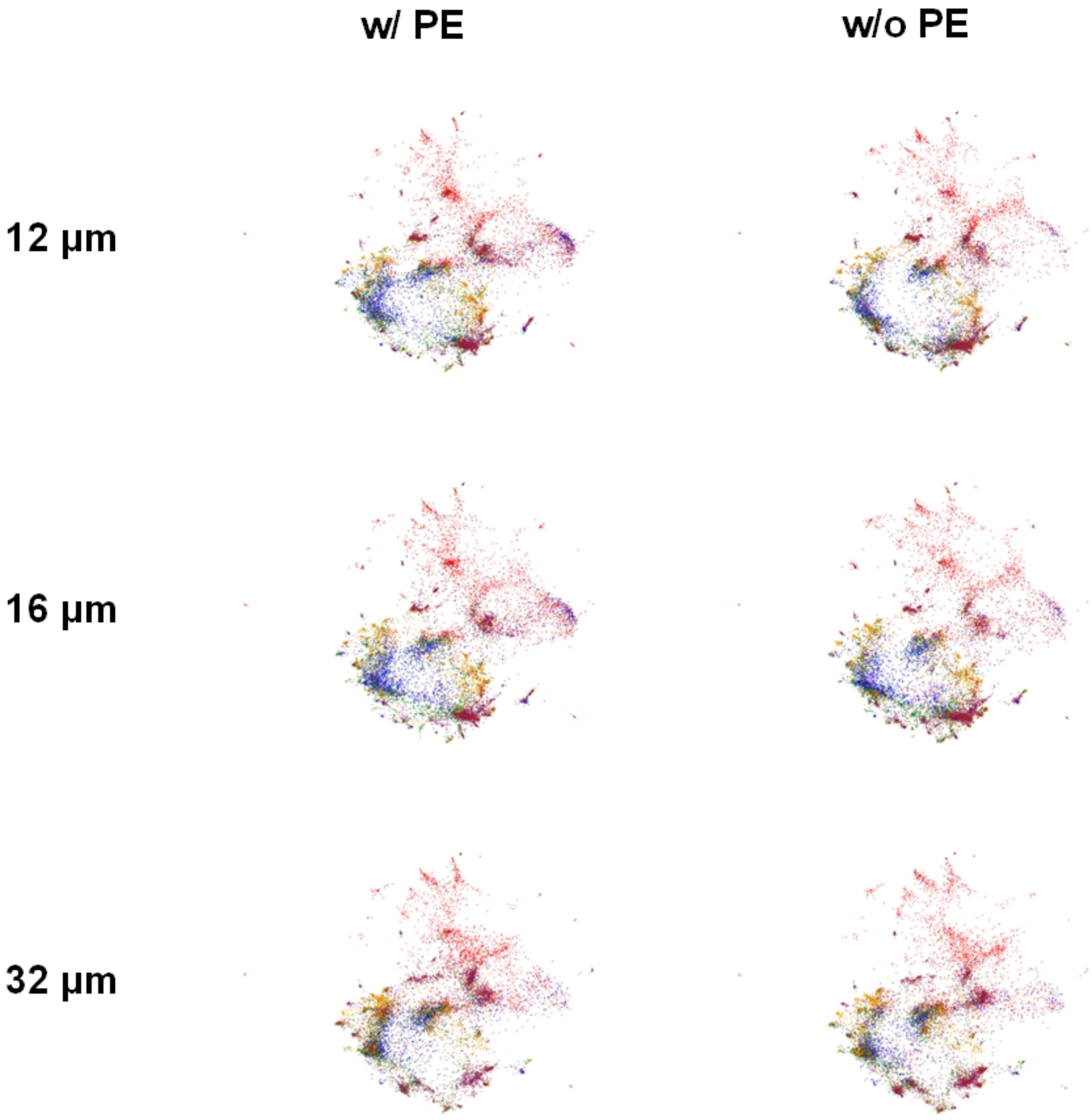


Figure 3. **Feature distributional.**

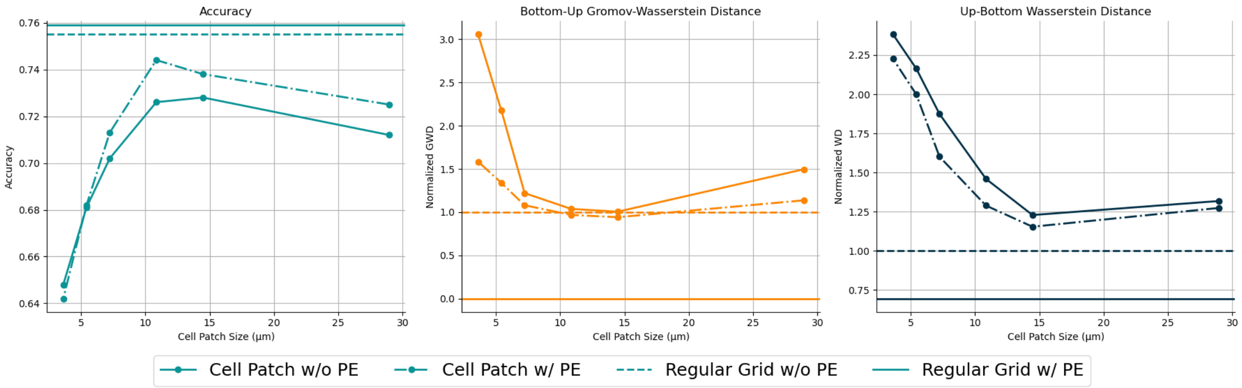


Figure 4. **Feature distributions distances.** Effects of cell patch size and positional embeddings are checked based on classification accuracy, bottom-up GWD $D_{bottom-up}$, and up-bottom WD $D_{up-bottom}$ on three subfigures.

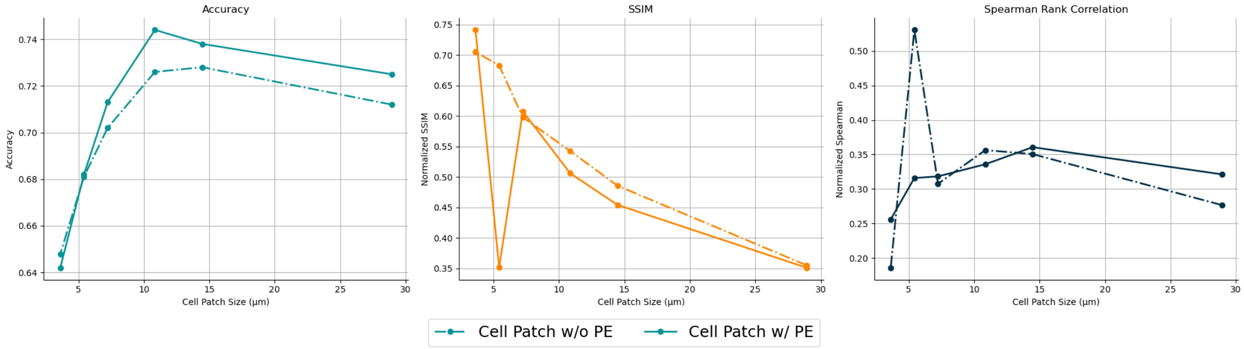


Figure 5. **Saliency map distances.** Effects of cell patch size and positional embeddings are checked based on classification accuracy, SSIM and SRC on three subfigures.

#### References


[1] Julius Adebayo, Justin Gilmer, Michael Muelly, Ian Goodfellow, Moritz Hardt, and Been Kim. Sanity checks for saliency maps. *Advances in neural information processing systems*, 31, 2018.

[11] Leland McInnes, John Healy, and James Melville. Umap: Uniform manifold approximation and projection for dimension reduction. *arXiv preprint arXiv:1802.03426*, 2018.
